# *pyRforest*: A comprehensive R package for genomic data analysis featuring scikit-learn Random Forests in R

**DOI:** 10.1101/2024.06.09.598161

**Authors:** Tyler Kolisnik, Faeze Keshavarz-Rahaghi, Rachel Purcell, Adam Smith, Olin Silander

**Affiliations:** School of Mathematical and Computational Sciences, Massey University, Auckland, New Zealand; Canada’s Michael Smith Genome Sciences Centre at BC Cancer, Vancouver, British Columbia, Canada; Department of Bioinformatics, University of British Columbia, Vancouver, British Columbia, Canada; Department of Surgery, University of Otago, Christchurch, New Zealand; The Liggins Institute, University of Auckland, Auckland, New Zealand

**Keywords:** Bioinformatics, Machine Learning, Biomarker Identification, Random Forest, Genomic Data Analysis

## Abstract

Random Forest models are widely used in genomic data analysis and can offer insights into complex biological mechanisms, particularly when features influence the target in interactive, non-linear, or non-additive ways. Currently, some of the most efficient Random Forest methods in terms of computational speed are implemented in Python. However, many biologists use R for genomic data analysis, as R offers a unified platform for performing additional statistical analysis and visualization. Here we present an R package, *pyRforest*, which integrates Python *scikit-learn* `RandomForestClassifier` algorithms into the R environment. *pyRforest* inherits the efficient memory management and parallelization of Python, and is optimized for classification tasks on large genomic datasets, such as those from RNA-seq. *pyRforest* offers several additional capabilities, including a novel rank-based permutation method for biomarker identification. This method can be used to estimate and visualize p-values for individual features, allowing the researcher to identify a subset of features for which there is robust statistical evidence of an effect. In addition, *pyRforest* includes methods for the calculation and visualization of SHapley ADditive Explanations (SHAP) values. Finally, *pyRforest* includes support for comprehensive downstream analysis for gene ontology and pathway enrichment. *pyRforest* thus improves the implementation and interpretability of Random Forest models for genomic data analysis by merging the strengths of Python with R. *pyRforest* can be downloaded at: https://www.github.com/tkolisnik/pyRforest with an associated vignette at https://github.com/tkolisnik/pyRforest/blob/main/vignettes/pyRforest-vignette.pdf.

## Introduction

The field of genomics has evolved over the past two decades, from studies primarily focused on individual genes to comprehensive analyses of entire genomes [1]. This shift has been facilitated by advances in sequencing technologies and bioinformatics tools, allowing researchers to generate and analyze vast quantities of genomic data [2]. To derive meaningful biological insights from these large datasets, advanced statistical methods, such as machine learning, are increasingly required.

Decision trees are a supervised learning method frequently used for classification. The values of different features are used to split data along branches to make classification predictions. Random Forest (RF) models are ensembles of random trees, i.e., decision trees where only a random subset of features are used at each split, known for their high predictive power and robustness to dataset noise for handling large datasets, such as those of genomic research [3]. RFs can mitigate overfitting through mechanisms specifically designed to introduce variability among the individual trees, such as bootstrapping and feature bagging. These properties make RF models particularly competitive for binary classification problems [4].

Python [5] is often used for implementing RF models due to having libraries designed with efficient memory handling and parallelization in mind. However, R [6] is commonly used as a computing environment by biologists, though implementations of RF models in R are slower and less optimized compared to Python [7]. Many RF implementations in R suffer from inefficient memory handling and lack parallelization strategies, which are crucial when processing large genomic datasets [7]. This creates a barrier for researchers who must often choose between the advanced machine learning capabilities of Python and the comprehensive genomic analysis tools available in R.

To address these limitations, we present *pyRforest*, an R package that integrates the *scikit-learn* `RandomForestClassifier` algorithm [8] implemented in Python into the R environment using the *reticulate* package [9]. *pyRforest* enables users familiar with R to leverage the machine learning strengths of Python without requiring any Python coding knowledge. This integration improves memory management and parallel processing, allowing the user to create RF models on larger genomic datasets with less RAM usage when compared to R, while enhancing the interpretability and biological relevance of RF models in genomic studies.

In addition, *pyRforest* offers several innovative features, including a novel rank-based permutation method for identifying significantly important features, by estimating and visualizing p-values for individual features. This allows researchers to prioritize a reduced list of biomarkers for further analysis, while speeding up computation. Additionally, *pyRforest* includes methods for calculating and visualizing SHapley ADditive Explanations (SHAP) values [10, 11] while also supporting comprehensive downstream analysis for gene ontology and pathway enrichment using *clusterProfiler* [12] and *g:Profiler* [13]. By merging the computational strengths of Python with the statistical and visualization capabilities of R, *pyRforest* addresses current limitations in genomic research workflows, contributing to the ongoing evolution towards more versatile and integrated bioinformatics tools and the need for more explainable artificial intelligence [14].

This paper introduces and outlines the capabilities of the *pyRforest* package for R. We first describe the development, use, and functionality of *pyRforest*. Next, we compare *pyRforest* to existing RF implementations, highlighting its computational advantages and innovative features. We then present a case study on colorectal cancer (CRC), demonstrating the utility of *pyRforest* in identifying key biomarkers and providing insights into their biological implications. Finally, we discuss the advantages and limitations of the package, as well as potential future directions for its development and application.

## Methods and Design

### R Package Development

*pyRforest* was created using the Rstudio IDE for R [15]. The *devtools* package [16] was used to streamline the code development, testing and documentation of the package. Github was used for version control and collaboration [17].

### Integration of R and Python

The *pyRforest* setup process supports the built-in *virtualenv* environments of *reticulate* [9] as well as *conda* [18] environments for Python package management. In our experience, *virtualenv* works best on Linux systems, and c*onda* is most suitable for Windows or Apple Arm64/M series Mac users.

For detailed setup instructions, including a step-by-step guide for configuring *virtualenv* or *Conda* with *reticulate*, we refer users to a vignette included with *pyRforest*. The vignette includes a demonstration of the capabilities of *pyRforest* with an example dataset. The development of *pyRforest* and all case studies were carried out on an M1 iMac with 16GB RAM. The package was also tested successfully on a Windows 10 x64 system with an Intel i5 processor, and Linux Ubuntu 20.04 LTS systems.

### Dataset Preparation and Hyperparameter Optimization

*pyRforest* includes functions for formatting and partitioning data into training, validation, and testing sets. *pyRforest* optionally supports multiple class weighting strategies to ensure balance of the target variable across the data partitions, which can be particularly important if the sample size is unbalanced with respect to the target class.

*pyRforest* optionally uses *scikit-learn* `BayesSearchCV` or `GridSearchCV*`* [8] to optimize RF hyperparameters, with default settings optimized for the genomic datasets presented in this paper. However, *pyRforest* allows users to customize hyperparameters, such as *‘*n_estimators*’* (number of decision trees), *‘*max_features*’* (maximum number of criteria for node splits), and *‘*max_depth*’* (maximum parent nodes on a decision tree), and others (see documentation for a full list) [8].

### Model Tuning and Evaluation

The model is initially trained using the training set, and tuned using the validation set, with final assessment scores calculated on the testing set. The default metric for assessing model performance is the Area under the Receiver Operating Characteristic (ROC) curve (AUC) score, which is a combined measure of sensitivity and specificity. However, the user can choose from a range of other metrics, including precision and accuracy. The validation test set allows the user to test for model overfitting via the training set. For example, if training results in overfitting, such models will often underperform on the validation set; this poorer performance can be used as an indication of overfitting, requiring further tuning or pruning of hyperparameters.

After the hyperparameter-tuning phase, the testing set is used to assess model performance. Importantly, the testing set is not used in model fitting or in tests for overfitting. *pyRforest* provides the user with a range of scoring metrics, such as accuracy, ROC-AUC score, sensitivity, specificity, and F1 score.

### Post-Hoc Feature Importance Significance Testing

*pyRforest* includes a procedure for identifying which features are significantly important to the predictive performance of the final model. The importance of each feature within the dataset is measured using the Gini importance score, and features are ranked from most to least important.

The statistical significance of each feature at each rank is determined using permutation to generate a null distribution of importance score profiles at each rank, to which the true importance scores of features at that rank are compared. This approach allows the calculation of p-values for each ranked feature by comparing the observed importance score to the null distribution of importance scores obtained under permutation for the corresponding rank. By default, *pyRforest* performs 1,000 permutations (although this is customizable). This permutation test allows users to identify a subset of features for which there is statistical evidence of their importance.

### Generating a Null Distribution of Importance Scores at Each Rank

The null distribution of importance scores for each rank is generated using the following steps:

1. Fit RF model; calculate importance scores for each feature; rank features from highest to lowest importance score.
2. For each permutation (repeat 1000 times (default)):
  i. Randomly permute the values of the target variable across samples.
  ii. Fit RF model.
  iii. Calculate importance scores and rank them from highest to lowest.
3. For each feature according to its importance rank:
  i. Obtain the null distribution of importance scores under permutation for that rank.
  ii. Determine the p-value: the proportion of null importance scores that are greater than the observed importance score.

The p-values for each feature capture the probability of obtaining an importance score as great or greater than the one observed at its rank under a null hypothesis of no association between features and target variable.

In comparison, in the most common approach to RF analysis, variable importance is assessed by the change in accuracy after the permutation of the value of a variable is obtained in an out-of-bag sample [3, 19]. Other permutation-based methods generally fall into two types: in one, class labels are swapped, and for each feature, the distribution of importance scores (frequently, Gini) is tested relative to the unpermuted dataset. For example, the *rfPermute* package [20] uses the mean decrease in Gini score for all permutations relative to the non-permuted data. In the other type, class labels are swapped, and for each feature, the distribution of the ranks is tested relative to the unpermuted dataset [21]. This contrasts to the case of *pyRforest*, in which the feature importance (Gini) at each rank is compared to the importance of the feature at that rank in the unpermuted dataset. More succinctly, *pyRforest* uses rank-based feature importance rather than feature-based importance, or feature-based *rank* importance.

The result is a list of features for which, given a chosen significance threshold (alpha), there is statistical evidence for non-zero importance to the predictive outcome of the final model. In a biological context, these features may be further studied as a list of potentially important biomarkers, or to gain insight into biological mechanisms.

*pyRforest* also offers plotting functions to visualize these features, leveraging the popular R package *ggplot2* [22]. These plots can aid in the interpretation of the significance assessment. These permutation and visualization steps allow the researcher to determine which features (for example, genes) found by the RF model are deemed significant and worthy of further research.

### SHAPley Additive exPlanations (SHAP)

SHAP values help explain the predictions of RF models, offering insight into when, why, and how specific features are important in determining class membership [10]. To facilitate this, *pyRforest* offers built-in functions for calculating and plotting SHAP values from the *`*shap.TreeExplainer*`* class within the *SHAP* package [11]. SHAP analysis can complement biological contextualization by providing a means to interpret the effects of individual features on model predictions.

### Biological Interpretation

Finally, when using *pyRforest* on genomic data, *pyRforest* facilitates biological interpretation of the produced ranked lists of significant genes, which are automatically prioritized and formatted for compatibility with downstream analytical tools. As shown in the examples within the vignette, the resultant data format is suitable for direct integration with *clusterProfiler* [12] and *g:Profiler* [13], powerful platforms for biological annotation and analysis. Specifically, these tools enable users to contextualize the statistically significant features reduced by *pyRforest* within biological pathways, functions, and processes.

## Comparisons with alternative implementations

We first benchmarked memory usage and compute time for three different RF methods: R *randomForest*; *scikit-learn* `RandomForestClassifier` directly in Python; and our *pyRforest* implementation of *scikit-learn `*RandomForestClassifier` in R which leverages *reticulate*. **Table 1** shows the results of this benchmarking. As expected, *pyRforest* performed similarly to a direct implementation of *scikit-learn* `RandomForestClassifier` with some additional overhead memory and run time inefficiencies due to the integration layer between R and Python. Both *pyRforest* and *scikit-learn* `RandomForestClassifier` vastly outperformed R *randomForest* on large datasets. While R *randomForest* excelled with very small datasets, the inefficiencies in memory and run time rapidly scaled with dataset size. On our genomic training dataset of size 58,678 features x 248 samples, average memory usage of R *randomForest* was over 5x that of *pyRforest*, and the run time average was over 10x as long.

**Table 1.** A benchmarking analysis comparing memory usage and run time of Random Forest model fitting across different datasets, using R’s *randomForest*, Python’s *scikit-learn* `RandomForestClassifier`, and the *pyRforest* implementation of *scikit-learn `*RandomForestClassifier` in R. In the case of RNA-seq data, the target metric was ROC-AUC; for the Iris and Income datasets, the metric was accuracy. Execution time and memory usage are the average of five runs. In all cases, five-fold cross validation was used. Dataset sizes and classification types are: Iris, 150 samples, 4 features, multiclass; Income, 30,162 samples, 14 features, two-class; Demo RNA-seq, 40 samples, 101 features, binary; CRC RNA-seq, 248 samples, 58,678 features, binary.

| Dataset | Trees *† | Memory Usage (MiB)‡ |  |  | Execution Time (seconds) |  |  |
| --- | --- | --- | --- | --- | --- | --- | --- |
|  |  | randomFo<br>rest | scikit-learn | pyRforest | randomFo<br>rest | scikit-learn | pyRforest |
| Iris | 50 | 25.2 | 0.39 | 996 | 0.10 | 0.58 | 0.39 |
| Iris | 100 | 43.1 | 0.41 | 1000 | 0.12 | 0.62 | 0.42 |
| Iris | 150 | 62.5 | 0.53 | 1010 | 0.14 | 0.64 | 0.58 |
| Income | 50 | 4310 | 26.8 | 732 | 24.9 | 2.69 | 124 |
| Income | 100 | 8040 | 38.3 | 837 | 49.8 | 4.61 | 254 |
| Income | 150 | 11760 | 50.3 | 1260 | 75.7 | 6.58 | 377 |
| Demo<br>RNAseq | 50 | 14.5 | 0.38 | 1180 | 0.15 | 0.65 | 0.54 |
| Demo<br>RNAseq | 100 | 19.8 | 0.42 | 1180 | 0.18 | 0.67 | 0.377 |
| Demo<br>RNAseq | 150 | 25.2 | 0.52 | 1178 | 0.213 | 0.84 | 0.51 |
| CRC RNAseq | 50 | 13160 | 38.8 | 2290 | 308 | 3.70 | 34.3 |
| CRC RNAseq | 100 | 13190 | 40.7 | 2400 | 566 | 4.33 | 63.4 |
| CRC RNAseq | 150 | 13218 | 42.5 | 2454 | 822 | 5.12 | 92.5 |

\* All parameters were held constant where possible
† Number of estimators in Python is equivalent to Number of trees in R
‡ Average Times and Average Memory Usage represent an average over 5 repetitions

We also compared the benchmarking and results of different feature identification approaches in **Suppl. Table 1** using the CRC RNA-seq dataset. This table illustrates a comparative analysis of feature identification approaches, evaluating the number of important features with non-zero importance scores found, run time, and peak memory usage during benchmarking. Feature importance results were assessed on the default s*cikit-learn* `RandomForestClassifier`, *pyRforest, scikit-learn*’s `inspection` module with 1000 permutations, the *randomForest* R package, and the *rfPermute* R package. As shown in this table, *pyRforest’s* approach to rank-based permutation feature importance testing offers a considerably faster approach to feature identification relative to *scikit-learn*’s `inspection`, a Python package that also employs permutation testing. In multiple attempts at testing the R packages that utilize permutation, *randomForest* [19] and *rfPermute* (which relies on R *randomForest*) [20], the memory usage became prohibitive, ultimately causing the process to crash. This comparative analysis demonstrated the nuanced capability of *pyRforest* for feature selection. Of the 58,678 features available in the CRC RNA-seq dataset, *pyRforest* identified 83 genomic features with significant importance. This is a considerably reduced set compared to the 1008 features that had non-zero importance in the default *scikit-learn* `RandomForestClassifier*`* model without any feature prioritization applied. In contrast, we found the *scikit-learn* `inspection` module to be overly conservative in identifying features, as it identified only a single feature and missed several features that are well-established to affect CRC localization [23, 24].

Several other advantages of *pyRforest* relative to the two other RF implementations in R are shown in **Suppl. Table 2**. For example, in contrast to other tools, *pyRforest* innately supports direct class weighting, k-fold cross-validation, and simple integration with cross-validation tools, streamlining the analysis process.

**Table 2.** K-fold cross-validation scoring metrics for the RF model, stratified by dataset splits (Validation, Testing) on the CRC dataset from our case study. The table presents key performance metrics including Accuracy, F1 score, Precision, Recall, and ROC-AUC.

| Scoring Metric | Validation Set | Testing Set |
| --- | --- | --- |
| Accuracy | 0.87 | 0.8 |
| F1 | 0.88 | 0.8 |
| Precision | 0.78 | 0.8 |
| Recall | 1.0 | 0.8 |
| ROC-AUC | 0.88 | 0.8 |

In summary, *pyRforest* bridges the computational and methodological divide between Python and R, providing users with advantages of both: the advanced machine learning capabilities and efficiencies of Python *scikit-learn* and the comprehensive analysis strengths of R. It introduces an innovative rank-based permutation method tailored for biomarker prioritization in genomic data analysis. Additionally, pyRforest simplifies GO analysis with easy integration into *clusterProfiler* and *g:Profiler*. This unique combination of features positions *pyRforest* as a strong tool for end-to-end genomic studies.

## Case Study

### RNA-seq Data Analysis in Colorectal Cancer

In this case study, we employed *pyRforest* to analyze a dataset comprising RNA-seq data from colorectal cancer samples obtained from the University of Otago, Christchurch, New Zealand analyzed in our previous study [23]. This dataset includes 308 patient samples, and encompasses a comprehensive range of 58,678 genomic features, including genes and long non-coding RNAs (lncRNAs). The primary focus of this analysis was to distinguish the cancer localization within the colorectum based on the binary outcome variable ‘side’ (left vs. right).

### Data Preparation and Model Training

The raw genomic data was mapped to the human genome (GRCh38) using *STAR* (v2.73a) [25] and TPM normalized to remove sequencing depth and gene length biases. It was then split into training (248 samples), validation (30 samples), and testing (30 samples) sets, ensuring a robust evaluation for the RF model developed using *pyRforest*. The validation and testing sets were balanced for outcome, and the training set was minimally unbalanced with 57% left-sided samples and 43% right-sided samples. The RF model was trained and tuned using the *pyRforest* exhaustive grid search, yielding optimal hyperparameters noted in **Suppl. Table 3**. The initial training of the model on the training set identified 1008 features with non-zero Gini importance scores, indicating a significant level of complexity and feature interaction within the data **(Suppl. Table 4)**.

### Model Performance

In our case study, the RF model exhibited strong performance across a range of scoring metrics, including accuracy, F1 score, precision, recall and ROC-AUC score, as seen in **Table 2**. Notably, the model achieved high accuracy rates of 0.87 and 0.80 for the validation and testing sets respectively, with both F1 and ROC-AUC scores closely mirroring these values at 0.88 and 0.80. The model demonstrated strong generalization capabilities on the unseen testing dataset, and consistently high performance at discriminating between the outcome classes.

### Identifying Significantly Important Features

The rank-based permutation approach of *pyRforest* identified 83 significantly important biomarkers of the total available set of 58,678 **(Figure 1)**. Each feature was assigned a p-value, providing a statistical basis for their inclusion in the downstream analyses **(Suppl. Table 5)**.

**Figure 1.**
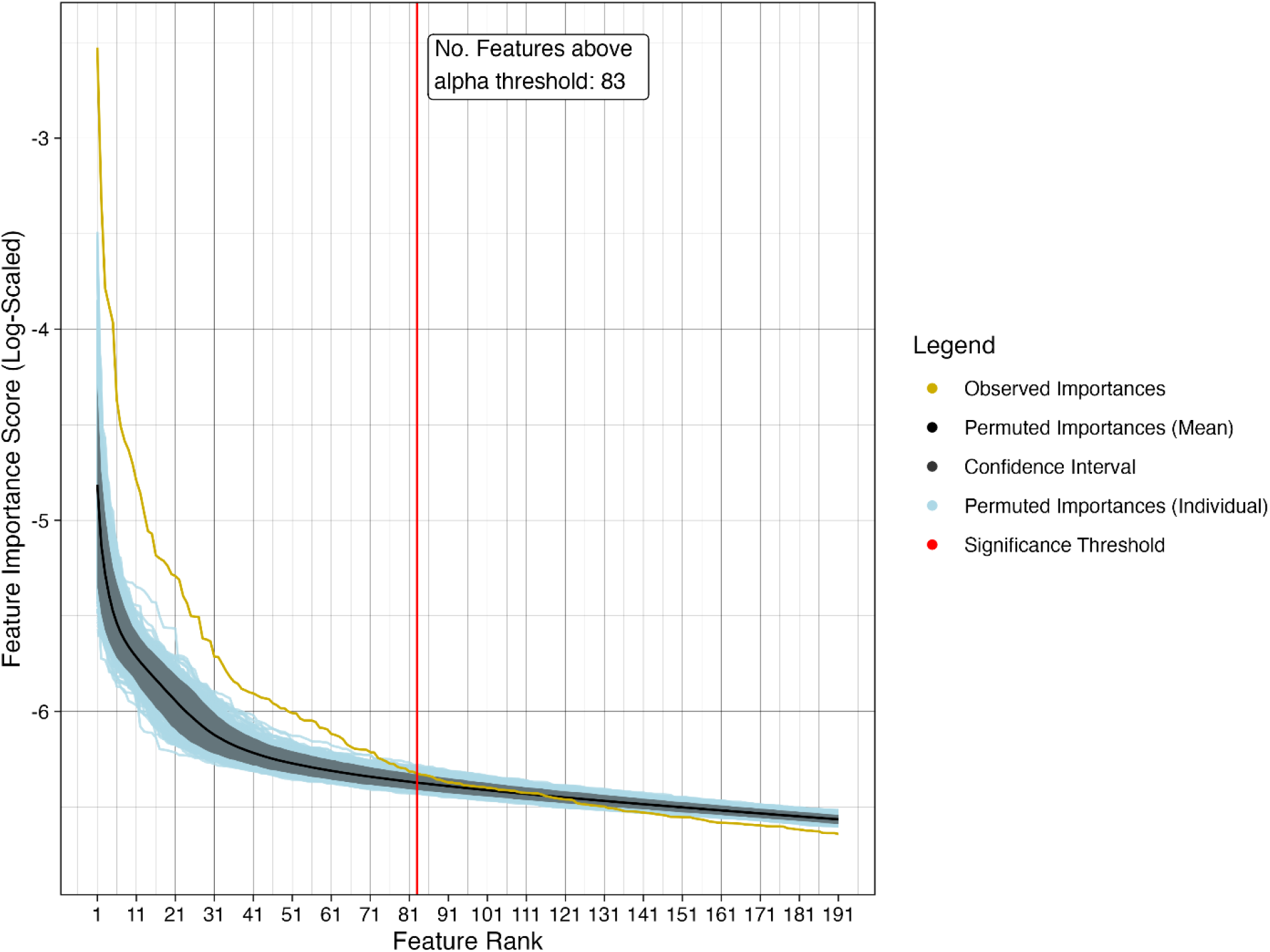
Feature importance plot showing individual and mean rank-based feature importance scores of the permuted data. Feature ranks are shown on the x-axis, with the importance score on the y-axis. The yellow line indicates the importance scores of the features at each rank in the unpermuted data. The blue lines indicate the importance scores at each rank for the permuted data sets, and the shaded gray region contains 95% of the ranked importance scores of the permuted data. The significance threshold was determined based on an alpha of 0.05 (i.e. the importance score of the ranked feature in the unpermuted is larger than the score in 95% of permuted data), and is indicated by the red vertical line.

Notably, the small nuclear protein *PRAC1* emerged as the top feature, indicating its potential role in the lateralization of CRC, which is a result consistent with many previous studies [23, 24].

*HOXB13*, and the lncRNA ENSG00000242407 were also identified as important features. *HOXB13* and this lncRNA colocalize with *PRAC1*: all are between 48.721 and 48.734 Mbp on chromosome 17. Their functional relationship to *PRAC1* is not clearcut. *HOXC4* and *HOXC6* were also identified; as developmental genes, these are functionally related to *HOXB13* and overlap on chromosome 12 between 53.991 and 54.017 Mbp. In addition, *QPRT* (tryptophan metabolism) and *WASF3* (cell movement and adhesion), were identified, both of known importance to CRC [26, 27]. Finally, *pyRforest* identified a second lncRNA, ENSG0000250829 and this has no identified function, suggesting new avenues for future research. *PRAC1* was the sole feature identified by the *scikit-learn `*inspection` feature identification approach **(Suppl.Table 4)**.

### SHAP Analysis

Calculation of SHAP values provides additional insights into the direction of the contribution (positive or negative) of these features to the target variable’s outcome within the RF model, here defined as associated with left-sidedness **(Figure 2)**. SHAP importance rankings are independent from *pyRforest* rankings. Specifically, *SHAP* analysis indicated that *PRAC1* and *HOXB13* contribute to the predisposition towards left-sided CRC. This observation is consistent with outcomes from other studies, highlighting the role of these biomarkers as being differentially expressed depending on spatial location of the disease [23, 24]. *HOXC4* and *HOXC6* SHAP directionality deviated from expected values from our previous study: the bimodal feature explanations of these genes indicate the relationship of these genes with outcome may be complex, and that their contribution is context dependent. In other words, depending on the expression level of other genes, these can contribute either positively or negatively toward left-sided CRC. SHAP analysis indicated that lncRNA ENSG0000250829 is the top feature contributing towards right-sided CRC, although the cross-shaped spread of the local feature explanation also indicates a complex, non-linear relationship with outcome. The inclusion of SHAP analysis enhances our understanding of the decisions of the predictive model, offering a clearer picture of how individual features can influence the output of the model.

**Figure 2.**
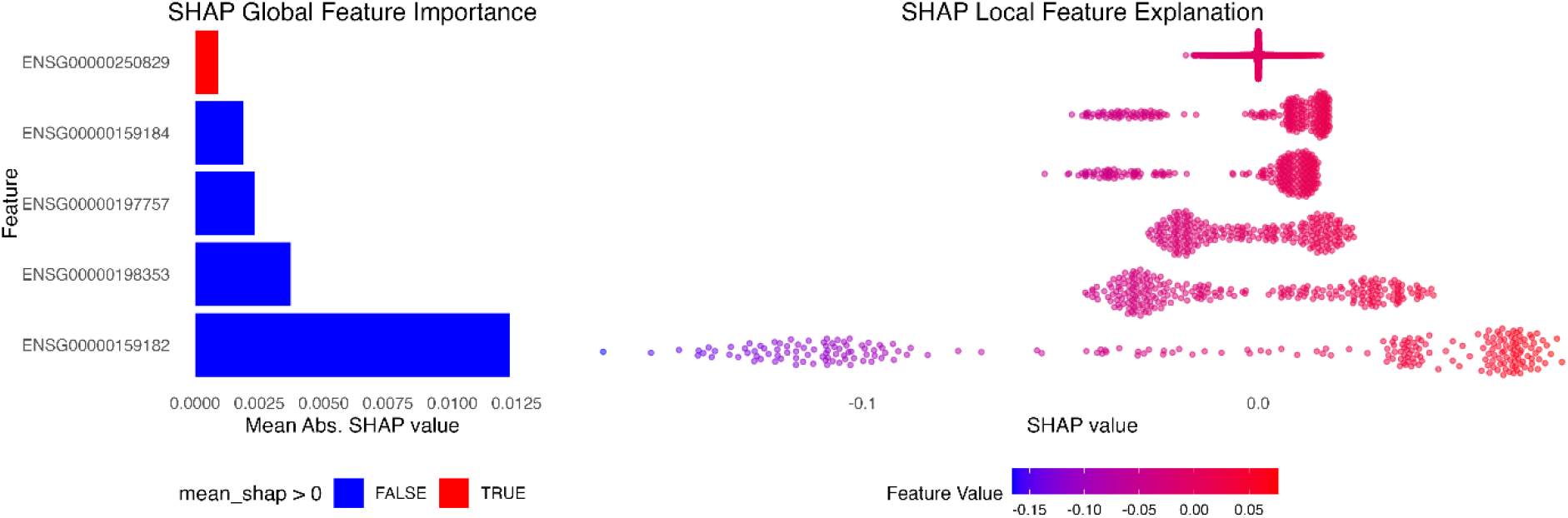
SHAPley Additive Explanation plots which illustrate the impact of individual features on model prediction. The left panel displays the global feature importance with the mean absolute SHAP values, where blue signifies features driving predictions towards ‘left-sided CRC’ and red signifies features driving predictions towards ‘right-sided CRC’. The right panel shows the distribution of individual SHAP values for each feature, reflecting their contribution to each prediction - negative SHAP values suggest a push towards ‘left-sided CRC’, whereas positive values indicate a push towards ‘right-sided CRC’. The color intensity corresponds to the feature value magnitude, with cooler colors representing lower values and warmer colors indicating higher values. From top to bottom, the ENSEMBL Gene IDs in the figure above correspond to: the novel lncRNA transcript *AC108865*.*1*; *HOXB13*; *HOXC6*; *HOXC4*; and *PRAC1*.

### clusterProfiler and g:Profiler Analysis

pyRforest returns a list of genes that can easily be used as input for the R packages *clusterProfiler* and *g:Profiler*. The *clusterProfiler* gene ontology (GO) enrichment analysis using the 83 identified features, revealed involvement in significant biological processes such as ‘anterior/posterior pattern specification’ and ‘cytolytic granule’ **(Figure 3)**. The top results from *g:Profiler* gene enrichment analysis also reinforce the findings that there is a potential link between prostate development processes and CRC anatomical side **(Figure 4)**. The enriched *GO* terms provide biological plausibility to the findings of the case study, indicating that the significant features identified by *pyRforest* are not only statistically relevant but also biologically grounded in the context of CRC lateralization.

**Figure 3.**
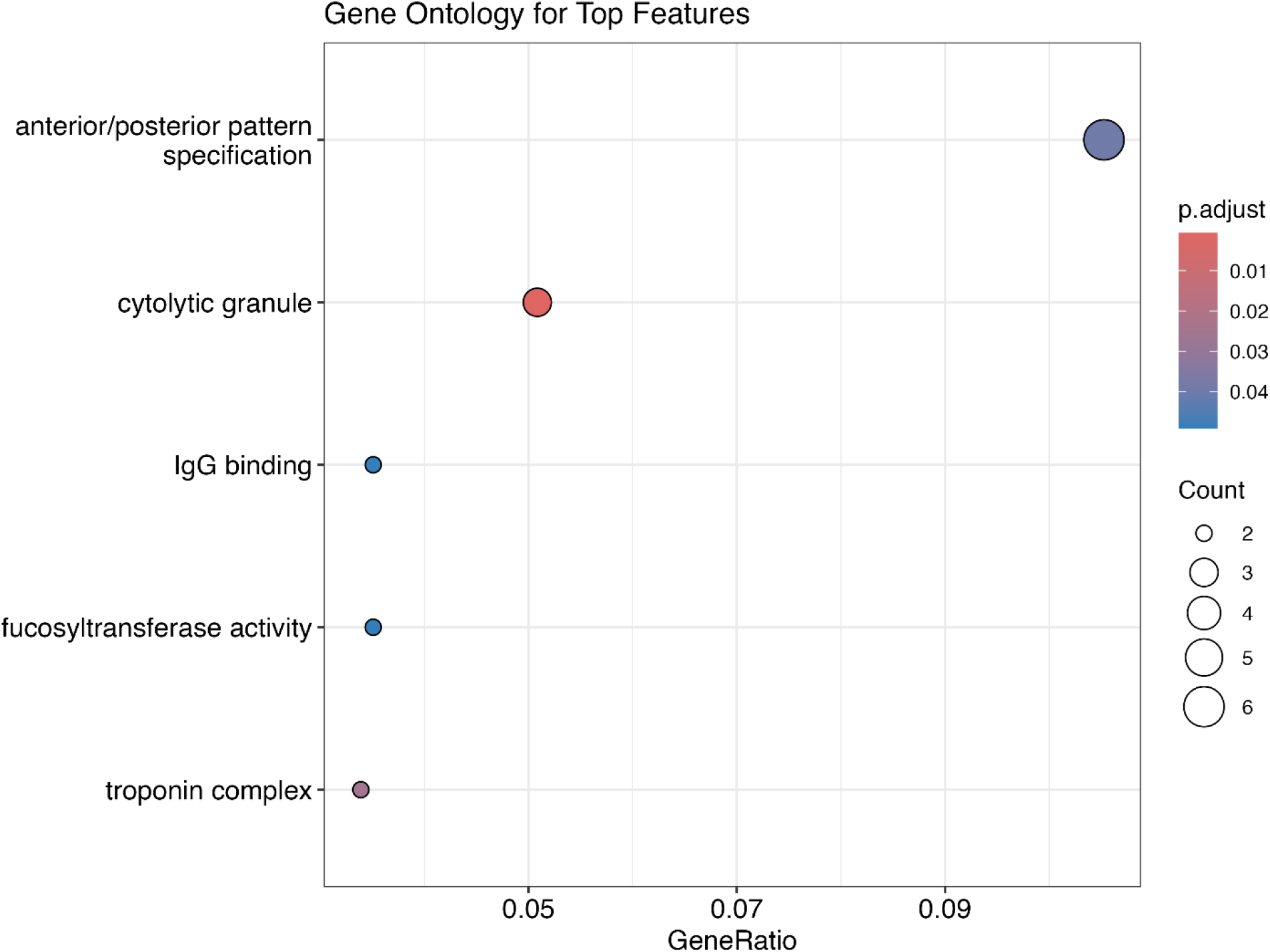
Gene Ontology enrichment analysis using clusterProfiler on the significant features identified by pyRforest. The x-axis represents the GeneRatio, the proportion of genes involved in the GO term relative to the total number of genes studied. The y-axis lists the GO terms associated with the identified features. Circle size indicates the gene count associated with each term, while the color gradient represents the adjusted p-value (p.adjust), with cooler (bluer) colors indicating less significance.

**Figure 4.**
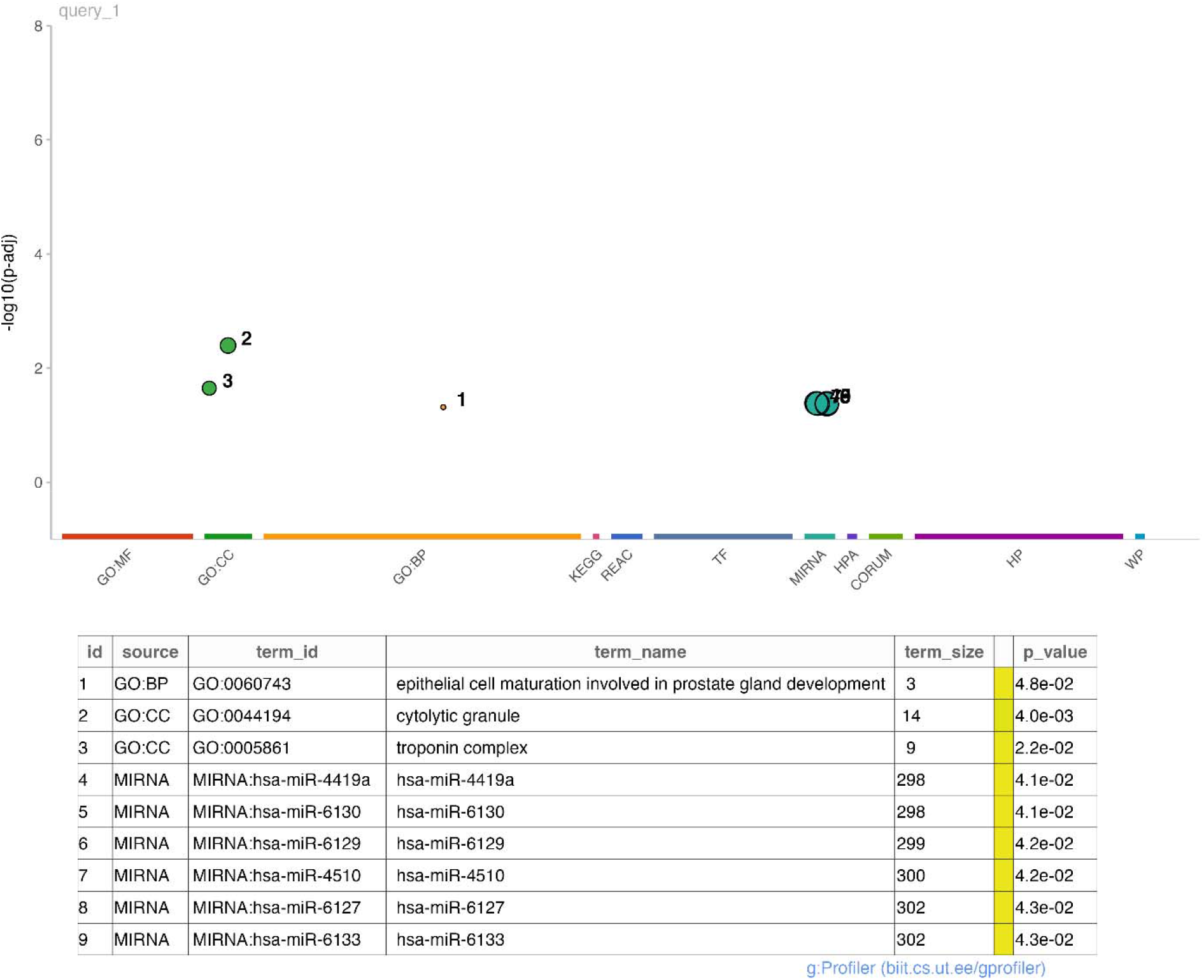
Enrichment analysis Manhattan plot from g:Profiler calculated on the 83 significant features found in our CRC case study. The x-axis categorizes enriched terms from various databases, and the y-axis shows their significance (-log10 p-value). Points represent terms, with notable significantly enriched terms named in the table below the plot.

## Discussion

*pyRforest* combines the computational power of Python and the statistical and genomic analysis capabilities of R into one package, addressing a crucial need in modern genomic research.

*pyRforest* offers computational speed and memory efficiency advantages over common R-based RF implementations **(Table 1)**. Moreover, *pyRforest* offers a range of extra utility functions for biomarker selection and analysis. These include (1) a rank-based permutation approach which allows the user to identify the features that have statistically significant importance scores, (2) integration SHAP for feature interpretation and (3) compatibility with *clusterProfiler* and *g:Profiler* for producing gene enrichment results which allow the user to explore the list of significantly important features in the context of external databases of known biological functions **(Figures 3 and 4)**.

A primary objective of analyzing genomic data with RF models is to discern important biomarkers. We present a comparison of *pyRforest* with two other packages which each differ in their approach to using permutation to identify important features. The ‘permutation_importance*’* function in the *scikit-learn* `inspection` module [8] independently permutes features, which is computationally intensive and destroys any associations among features. In contrast, *pyRforest* permutes only the response variable, thereby leaving the relationships among features intact. This approach offers advantages in computational speed, and we consider this approach to yield a more appropriate null model, given the extent of the known correlations and gene-gene interactions present within real genomic data. Another R package, *rfPermute* [20], acts as a wrapper for the *randomForest* R package and, like *pyRforest*, permutes only the target variable rather than the features. However, it compares the importance of each feature to its own distribution of importances scores under a permuted target variable. In contrast, *pyRforest* compares each observed feature importance to the distribution of permuted importance scores at its rank, rather than to the distribution of permuted importance scores of the feature itself. The null distribution for the most important feature (rank 1) is the scores obtained at rank 1 under permutation, regardless of the possibility of the identity of the specific feature changing with each permutation.

We consider that these two properties, permuting the target variable and not the features, and using rank-based importance distributions to offer a more appropriate null model and basis for comparing observed importance scores, and, ultimately, identifying a reduced set of potentially important features.

The approach of *pyRforest* also leads to two key effects that differentiate it from other rank-based methods, such as that of the previously mentioned *rfPermute* [20], and the approach implemented in Altmann et al. [20], which determines feature significance by comparing the rank of a feature in the test data to the distribution of ranks in permuted data. First, *pyRforest* can more effectively select important variables in datasets where a large proportion of the variables are predictive. For example, in a dataset with 100 variables, a variable’s importance rank in permuted data will typically center around 50 (with a standard deviation of approximately 7 under a Poisson distribution). However, if 50 or more variables are predictive, traditional methods may not detect any variable as significantly important compared to its average rank in the permuted data. In contrast, *pyRforest* may still identify the 50^th^ variable as important if its observed importance is significantly greater than in the permuted data. Second, *pyRforest* is more attuned to feature importances taking extreme values, especially in permuted datasets. Such values may be uncommon, but they are often crucial for identifying important features in an RF model. Feature-rank based methods can obscure these outliers, as they rely on non-parametric comparisons against a null distribution, potentially missing important features with high importance.

In addition, the *pyRforest* rank-based approach forgoes the need to permute and evaluate each feature individually, and this generally offers improved speed compared to feature-based approaches.

Despite its innovative contributions, the *pyRforest* R package comes with limitations worth noting. The primary challenge is the computational time associated with the permutation process, although benchmarking results indicate that *pyRforest* is significantly faster than the widely used package *scikit-learn* `inspection`. One potential enhancement could be to reduce computation time by fitting a theoretical probability distribution to an empirical distribution derived from a smaller subset of permutations [21]. Additionally, *pyRforest* is currently tailored specifically for classification problems, but extending its capabilities to support regression analysis would further enhance its versatility and allow it to be applied to additional machine learning problems.

In summary, despite these limitations, we have demonstrated the utility of *pyRforest* as a powerful analytical tool. In our case study on a CRC dataset, *pyRforest* identified 83 significant biomarkers and facilitated the exploration of gene ontology, SHAP values, and gene set enrichment, far surpassing the 15 biomarkers reported in the original study [23]. The integration with SHAP values allowed us to uncover new insights into how each feature influences cancer localization, highlighting *pyRforest’s* ability to uncover deeper biological insights. This package offers researchers a robust framework for biomarker discovery. Furthermore, our rank-based permutation approach was also successfully applied in a study by Keshavarz-Ragahi et al. *(includes Kolisnik)* [28] identifying key features associated with p53 activity, which enhances our understanding of TP53-related transcriptional signatures across various cancer types.

## Conclusion

By integrating Python’s efficient `RandomForestClassifier` algorithm, *pyRforest* enables researchers to leverage computationally efficient machine learning approaches while staying in the R ecosystem. *pyRforest* offers a suite of tools for fitting, interpreting, and contextualizing RF models specifically for genomic studies. Its rank-based permutation approach for biomarker identification, alongside SHAP analysis, and integration with gene ontology tools aims to improve the interpretability and biological interpretation of RF models. These features make *pyRforest* a valuable resource for conducting and interpreting genome-scale RF studies.

Furthermore, performance improvements in the *scikit-learn* package can be easily integrated into *pyRforest* ensuring that the package remains up to date without requiring extensive code modifications. We encourage the research community to utilize, contribute to, and build upon *pyRforest*.

### Key Points

□ We developed *pyRforest*, an R package which offers seamless integration of *scikit-learn* `RandomForestClassifier`, `BayesSearchCV`, `GridSearchCV`, and `shap.TreeExplainer` modules into the R ecosystem via *reticulate*.
□ *pyRforest* introduces a rank-based feature importance permutation test designed for biomarker identification, which prioritizes and assigns statistical importance to RF model features based on Gini Importance scores.
□ *pyRforest* facilitates the exploration of the underlying biological mechanisms of RF models through *ggplot2* visualizations, and downstream integration with *clusterProfiler* and *g:Profiler*.
□ We demonstrate the effectiveness and utility of *pyRforest* in a colorectal cancer case study, where it identifies key biomarkers previously known to the literature and provides insights into the underlying biological mechanisms of the RF model.
□ Download, collaborate, make suggestions or report bugs at: https://www.github.com/tkolisnik/pyRforest.

## Supplementary Material

Supplementary data can be found online at *Briefings in Functional Genomics*.

## Author Statements

## Acknowledgements

We would like to acknowledge the Department of Surgery at the University of Otago for providing the patient samples and RNA-seq data used in the case study.

## Authors’ contributions

TK conceived the project, wrote the code and documentation for the R package, performed testing, analysis of results and figure generation, and wrote the first draft and contributed to all versions of the manuscript. FK assisted with testing of the *pyRforest* package and manuscript proof-reading. RP provided data and input on methodology and analysis. AS guided the development of statistical methods and validation, manuscript proof-reading, and guided presentation of results. OKS provided support with development, troubleshooting, statistics, and critical analysis of results, and contributed to the manuscript. All authors have read and approved this manuscript.

## Data Availability

Raw Sequence Reads are available at SRA BioProject ID: PRJNA788974 [29].

## Code Availability

The R package *pyRforest* used in these analyses is available at: www.github.com/tkolisnik/pyRforest [17].

## Ethics Approval and consent to participate

This study was approved by the University of Otago, New Zealand, Human Research Ethics Committee (approval number: H16/037). Informed consent was obtained from all subjects and/or their legal guardians. All experiments were performed in accordance with relevant ethics guidelines and the Declaration of Helsinki.

