## Supplementary Table 1 for "*pyRforest*: A comprehensive R package for genomic data analysis featuring scikit-learn Random Forests in R"

**Supplementary Table 1.** A comparative analysis of feature identification approaches, evaluating the number of features with non-zero importance, run time, and peak memory usage during benchmarking. Feature importance results were assessed on the default s*cikit-learn* `RandomForestClassifier`, *pyRforest,* *scikit-learn `*inspection` ‘permutation_importance’ with 1000 permutations, the *randomForest* R package, and the *rfPermute* R package.

| **Method** | **N Features with Non-Zero Importance** | **Run Time (hours)** | **Peak Memory Usage (16GB Max)** |
| --- | --- | --- | --- |
| Default s*cikit-learn* `RandomForestClassifier` | 1008 | 8 h | 10GB |
| *pyRforest* | 83 | 6 h | 8GB |
| *scikit-learn* `inspection` ‘permutation_importance’ (1000 feature-based permutations) | 1 | 95 h | 16GB |
| *randomForest* R package | NA | NA | 16GB |
| *rfPermute* R package | NA | NA | 16GB |
