## Supplementary Table 2 for "*pyRforest*: A comprehensive R package for genomic data analysis featuring scikit-learn Random Forests in R"

**Supplementary Table 2.** Feature and Capability Comparison of RandomForest Implementations for Genomic Data Analysis.

| **Feature/**  **Capability** | **Scikit-learn `RandomForestClassifier`** | **pyRforest** | **R randomForest** | **Rfpermute** |
| --- | --- | --- | --- | --- |
| Direct Class Weighting | Yes (via class_weight parameter) | Yes (via class_weight parameter) | No | Yes, via a preprocessing function |
| Parallel Processing Support | Yes (via n_jobs parameter) | Yes (via n_jobs parameter) | Requires manual setup or additional packages | Yes but memory inefficient, poor batching |
| Automatic Out-of-Bag (OOB) Error Estimate | Yes (via oob_score parameter) | Yes (via oob_score parameter) | Yes | Yes |
| Handling Missing Values | No (requires preprocessing) | Yes - preprocesses missing values | Yes (automatic handling with na.action) | Yes |
| Cross-Validation Integration | Easy integration with *scikit-learn* based cross-validation tools | k-fold cross-validation integrated | Requires manual setup or additional packages | Requires manual setup or additional packages |
| Grid Search for Hyperparameter Tuning | Direct integration with *scikit-learn `GridSearchCV`* | Direct integration with *scikit-learn `GridSearchCV`* | No. Requires extensive manual setup or additional packages | No. Requires extensive manual setup or additional packages |
| Built-in Feature Importance Evaluation | Yes (attribute feature_importances_) | Yes provides base model feature importance scores and permuted feature importance scores | Yes, with options for different importance types | Yes, with options for different importance types |
| Feature identification | Requires additional modules such as sklearn.inspection | Yes: null-distribution permutation rank based feature identification | Requires manual setup or additional packages | Yes: null-distribution permutation feature based feature identification |
| Extensive Ecosystem Compatibility | High (Python data science stack) | Extremely High - Compatible with both Python ecosystems for machine learning and R ecosystems for bioinformatics | Moderate (R ecosystem, less extensive for machine learning, but more extensive for bioinformatics) | Moderate (R ecosystem, less extensive for machine learning, but more extensive for bioinformatics) |
