## Supplementary Table 3 for "*pyRforest*: A comprehensive R package for genomic data analysis featuring scikit-learn Random Forests in R"

**Supplementary Table 3.** The top-scoring model hyperparameters as optimized by the *pyRforest* `tune_and_train_rf_model*`* function which makes use of *scikit-learn* `GridSearchCV*`*.

| **Parameter** | **Setting** |
| --- | --- |
| Bootstrap | TRUE |
| class_weight | balanced |
| criterion | gini |
| max_depth | 8 |
| max_features | 0.2 |
| min_samples_leaf | 4 |
| min_samples_split | 2 |
| n_estimators (# trees) | 100 |
| warm_start | FALSE |
